# Predicting High-Resolution Spatial and Spectral Features in Mass Spectrometry Imaging with Machine Learning and Multimodal Data Fusion

**DOI:** 10.1101/2025.02.19.639070

**Authors:** Md Inzamam Ul Haque, Ramakrishnan Kannan, Jacob D. Hinkle, Sylwia A. Stopka, Nathalie Y. R. Agar, Olga S. Ovchinnikova, Debangshu Mukherjee

## Abstract

Recent advancements in molecular Mass Spectrometry Imaging (MSI) have sparked interest in integrating high spatial resolution methods with molecular mass-spectrometry-based chemical imaging. Fusion-based algorithms have proven effective in generating high spatial-resolution molecular mass spectra. However, a significant challenge stems from the differing physical mechanisms underlying image generation and data upsampling techniques, potentially leading to discrepancies in integrated information channels. Integrating physical constraints into data processing workflows is essential to tackle this issue. In this study, we propose an innovative approach that merges data from Fourier transform ion cyclotron resonance (FTICR), time-of-flight matrix-assisted laser desorption/ionization (MALDI-ToF), and time-of-flight secondary ion mass spectrometry (ToF-SIMS) imaging techniques. By leveraging FT-ICR’s unparalleled spectral resolution and ToF-SIMS’s exceptional spatial resolution, we achieve submicron spatial resolution, enabling the observation of intact molecular species with remarkable spectral precision. Canonical correlation analysis is employed to incorporate physical constraints. Through sophisticated image processing and machine learning techniques, the results of this fusion hold significant promise for advancing our comprehension of complex systems and unveiling concealed molecular intricacies.

## 1 Introduction

In recent years, significant attention has been directed toward advancing molecular Mass Spectrometry Imaging (MSI) platforms at the nanoscale to achieve subcellular resolutions. Concurrently, researchers have been exploring innovative approaches to combine molecular mass-spectrometry-based chemical imaging with other imaging modalities to provide additional sample information that could be used to improve spatial information collected via MSI. These techniques encompass a diverse array of methods, ranging from scanning probes [1, 2], optical microscopy [3, 4] to electron and ion systems [5], each contributing its unique strengths to the evolving field of molecular imaging.

One notable achievement in this pursuit is the generation of high spatial resolution molecular mass spectra. This accomplishment has been made possible through the application of fusion-based algorithms [6–8], which combine data from various sources and modalities, creating a comprehensive molecular profile. However, despite their promise, these approaches encounter a fundamental challenge stemming from the disparate physical mechanisms underlying image generation for MSI and the techniques employed for data upsampling.

This divergence in underlying mechanisms poses a critical issue, leading to a potential mismatch between the integrated information channels. These algorithms rely on the correlation assumption between the various information channels but lack constraints based on known relationships. Consequently, the output of such algorithms is susceptible to reconstruction errors [9], undermining the accuracy and reliability of the generated molecular mass spectra.

Integrating physical constraints into data processing workflows is imperative to address this challenge and unlock the full potential of combining multiple information channels for scientific applications. By infusing these constraints, researchers can bridge the gap between the diverse imaging modalities and enhance the fidelity of molecular data integration, ultimately paving the way for more robust and accurate molecular analysis techniques in the future.

In this study, we explored an innovative approach that merges data from three distinct MSI imaging techniques: Fourier transform ion cyclotron resonance (FTICR), time-of-flight matrix-assisted laser desorption/ionization (MALDI-ToF), and time-of-flight secondary ion mass spectrometry (ToF-SIMS). By combining the exceptional spectral resolution of FT-ICR with the outstanding spatial resolution of ToF-SIMS, the goal was to achieve submicron spatial resolution, enabling us to observe intact molecular species with high spectral precision. This fusion of imaging modalities promises to advance our understanding of complex systems and uncover hidden molecular details through state-of-the-art image processing and machine-learning techniques.

## 2 Methods

The workflow begins with applying sparse Non-Negative Matrix Factorization (NMF) to all three modalities. These three-dimensional datasets, which contain spectral signatures at each coordinate point as the third dimension, are stored in a space-efficient sparse format. During the NMF process, the MALDI-TOF and ToF-SIMS datasets are resampled to align with the FT-ICR spectral resolution, bringing all three modalities to the highest spectral resolution. Next, a Python-based machine learning workflow is employed to process and co-register these combined datasets. Upon successful registration, canonical correlation analysis (CCA) enhances low-resolution FT-ICR MSI images to match the high-resolution ToF-SIMS resolution by learning the correlation between these modalities. Similar work has been done utilizing MALDI-TOF and ToF-SIMS in [10]. Here, we have utilized FT-ICR to get an even better spectral resolution.

### 2.1 Datasets

For the FT-ICR dataset, nineteen tissue sections of 12-m thickness were prepared and analyzed using a 9.4 Tesla SolariX mass spectrometer (Bruker Daltonics, Billerica, MA) in the positive ion mode with a spatial resolution of 100 *µ*m. The MSI data were exported from SCiLS lab 2020a (Bruker, Bremen, Germany) in the standardized format imzML and converted to the HDF5 format for machine learning analysis. ToF-SIMS and MALDI-ToF datasets were acquired in Harvard Medical School (**TODO: Please explain briefly MALDI-ToF, and ToF-SIMS data**). Again, these two datasets are converted to HDF5 format for computational analysis.

### 2.2 Sparse NMF

All three datasets ranged from a couple hundred gigabytes to a couple of terabytes. Hence, the dimension reduction technique was necessary to run the computational analysis successfully. NMF was chosen for its non-negativity. In this case, why sparse NMF is used instead of traditional NMF is described below. We take the same approach of performing Sparse NMF for FT-ICR, ToF-SIMS, and MALDI-TOF data. Procedures and results of the sparse NMF for all three datasets are described below.

#### 2.2.1 FT-ICR

The FT-ICR dataset was collected over multiple slices of a mouse brain. The dataset itself is three-dimensional, as it’s collected over a spatial region of several millimeters, and at each collected point, there is a spectral signature associated with the data. However, this data is stored in a sparse format. As shown in Figure 1(a), spectra are not collected from every location. Black regions are the regions from which no spectra were collected. Thus, even though the spatial dimensions of the image in Figure 1(a) are 238 × 672 = 159936, spectra exist only from 72078 positions, or around 45% of positions.

**Figure 1.**
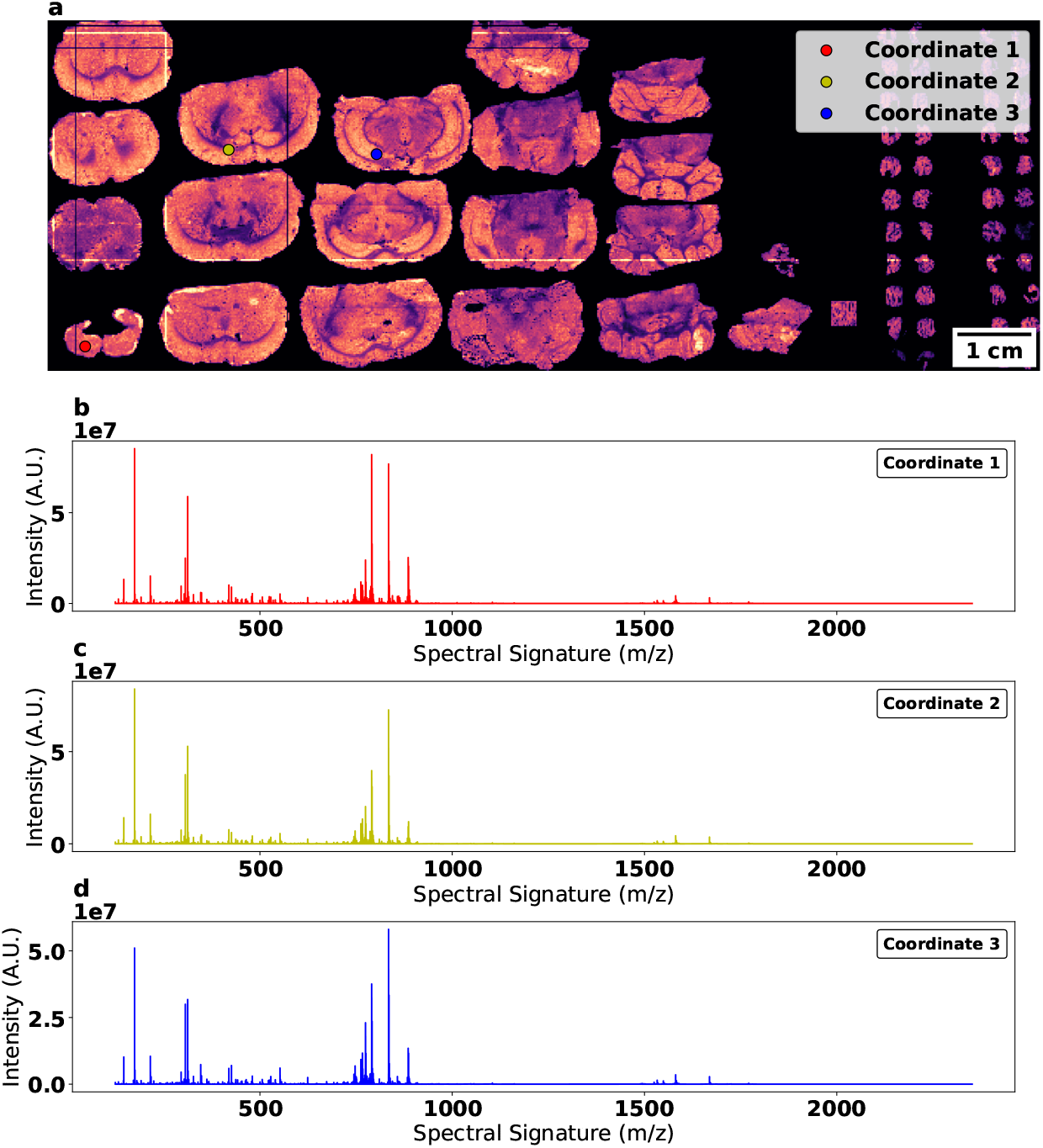
Spectral Components of FT-ICR. **(a)** Summed spectra from the FT-ICR data collected, with the individual mouse brain slices visible. **(b)** Spectra collected at point marked as Dataset 1. **(c)** Spectra collected at point marked as Dataset 2. **(d)** Spectra collected at the point marked as Dataset 3.

Spectra from three different locations, marked by points in Figure 1(a), are shown in Figure 1(b)-(d). Additionally, the data is collected and stored in a sparse format for each spectrum to save data size. This can be understood from looking at the spectra in Figure 1(b)-(d), where many of the intensities for higher m/z values, especially for those above 1000, are zero. Thus, comparison becomes highly challenging to such datasets - as each spectra can and will often have a unique set of m/z values. One way around this is to fill in the missing values with zeros and generate a filled-in three-dimensional dataset, where two dimensions are spatial and the third is spectral. However, such a dataset rapidly becomes extremely large and often cannot fit in memory. For example, when stored in the sparse format as just described, the dataset in question is around 5GB, while the filled data is over a hundred times larger, with most of the increase coming in from filling in the missing spectral values. Such a large dataset is almost impossible to fit in memory and extremely slow to analyze.

The solution to this impasse is to use Sparse NMF. To do this, we have used the Tensorly Python package. We start by finding the minimum distance between *m/z* points to scale all the unique *m/z* data at every spatial position. Then, we generate a three-column vector, where the first two columns refer to the *x* and *y* spatial positions, and the third column refers to the scaled *m/z* points, where the scaling factor was found in the previous step. We convert all the (*x, y*) spatial locations from the first two columns into a position list called z. This is obtained as *z* = *x* + (*y* × *y*_*max*_). We will use this as the proxy for spatial positions. Then, we load the *z* values and the spectral values into scipy.sparse.coo matrix. We then find all the unique components of *z*, which should be **72078**, the total number of spatial positions present in the data, and retain only the unique values in the coo matrix. We do the same for the spectral values. This ultimately results in a sparse matrix of 72078 × 9711 components, only 1.4 GB. Tensorly can natively handle sparse matrices through its’ sparse backend, which takes in a sparse coo matrix and outputs a sparse tensorly matrix **A**. Subsequently, tensorly’s sparse non-negative-parafac is used to decompose the sparse tensor (**A**) to the **W** and **H** matrices. We chose 40 components for the decomposition, so while **A** is 72078 × 9711 rank matrix, **W** is a 72078 × 40 rank matrix, and **H** is a 9711 × 40 rank matrix.

Supplementary Figure S1 shows two NMF components and their corresponding spectra computed using the sparse NMF process discussed above. Supplementary Figure S2 shows summed spatial and spectral maps of the 40 NMF components.

#### 2.2.2 ToF-SIMS

Like the FT-ICR data, the ToF-SIMS data is a three-dimensional dataset with a spectral signature associated at each spatial point. This data is also stored in a sparse format. Again, as seen previously with the FT-ICR data, the ToF-SIMS spectra are not collected from every location. We used the steps taken with FT-ICR data for this dataset to perform sparse NMF with the Tensorly package. The steps resulted in a sparse matrix of 2559964 × 769 components, 6.3 GB in size for the ToF-SIMS data. One crucial point to be noted here is that we used the *m/z* values from the FT-ICR measurements here for mainly two reasons. First, they help compare the final NMF results together for later steps. Second, we wanted to retain the highest spatial and spectral resolution as our final result, and FT-ICR data has the highest spectral resolution. As for the result of the sparse NMF done on ToF-SIMS data, we show two similar figures previously shown with FT-ICR data. Supplementary Figure S3 shows two NMF components and their corresponding spectra. Supplementary Figure S4 shows summed spatial and spectral maps of the 40 NMF components.

#### 2.2.3 MALDI-ToF

MALDI-TOF is also a three-dimensional dataset and is stored in a sparse format. The spatial dimensions of the image are 2329 *×* 5054 = 11770766; spectra exist from 2030712 positions or only 17.25% of positions. For this dataset, we used the same steps as discussed previously to perform sparse NMF with the Tensorly package. The steps resulted in a sparse matrix of 2030712 *×* 596787 components, 11 TB in size for the MALDI-TOF data. Again, we used the *m/z* values from the FT-ICR measurements. We chose 40 components for the decomposition, so while the sparse matrix **A** is 2030712 *×* 596787 rank matrix, **W** is a 2030712 *×* 40 rank matrix, and **H** is a 596787 *×* 40 rank matrix. Supplementary Figure S5 shows two NMF components and their corresponding. Supplementary Figure S6 shows summed spatial and spectral maps of the 40 NMF components.

### 2.3 ToF-SIMS grid removal

During NMF, grid artifacts were seen for ToF-SIMS data as seen in Figure S3(a) and Figure S4(a). Due to scanning issues, we can see 80×80 square tiles in the ToF-SIMS summed spectra image in Figure 2(a). Since the ToF-SIMS is a three-dimensional dataset with spectral components, this image is produced by summing up the spectral dimension, resulting in a two-dimensional image of 2000×1280, with 400 square tiles of 80×80. As we can see from the image, these tiles have a similar function, which act as an artifact during the sparse NMF and CCA process. We adopt the following workflow to eliminate this grid artifact and produce better results.

**Figure 2.**
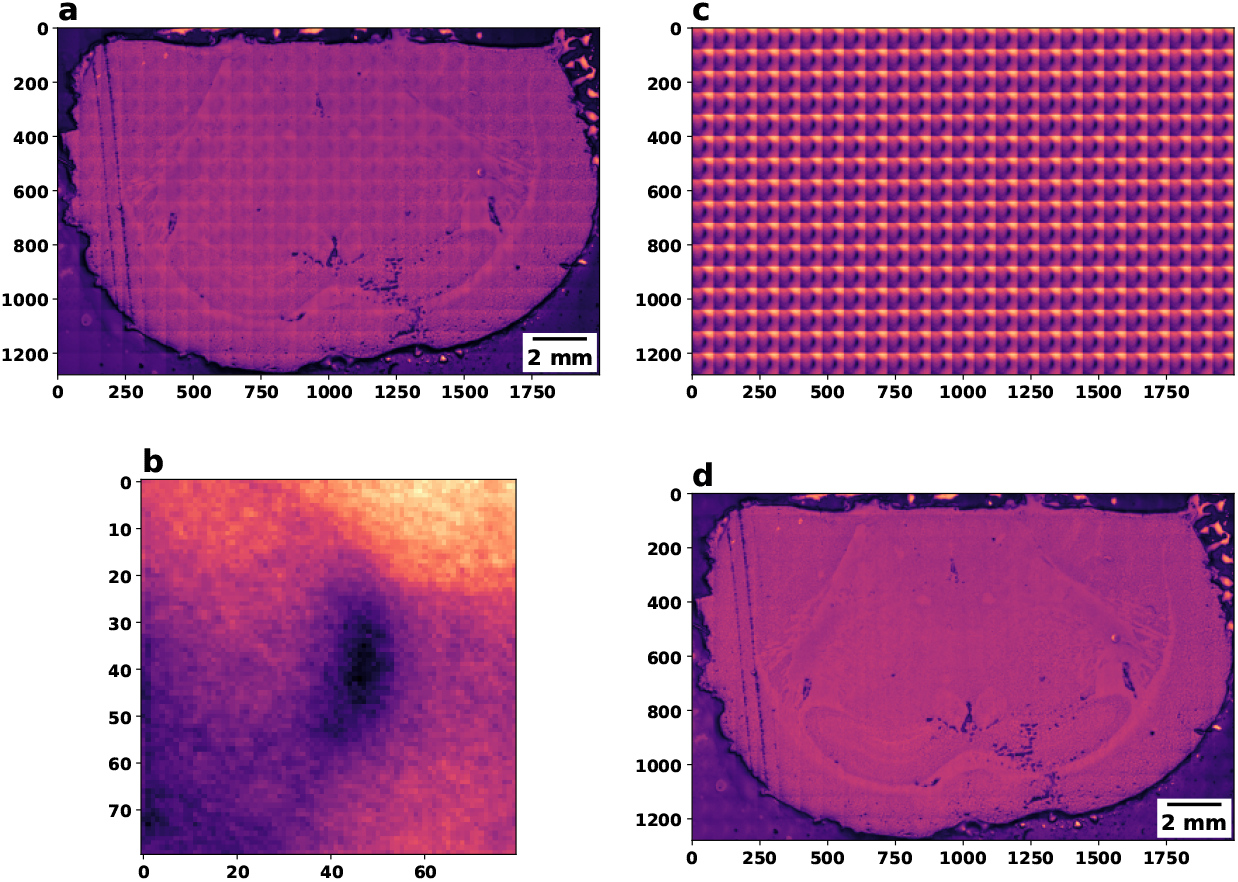
Grid artifact removal from ToF-SIMS dataset. **(a)** Original image - 2D spatial image by summing along the spectral dimension of the ToF-SIMS dataset. **(b)** arithmetic mean of the 400 tiles from **(a)** of size 80×80. **(c)** stacking the arithmetic mean tile to get a 2D image of the same size as the original image. **(d)** Resultant image after grid removal.

We get the 400 tiles from the image and take the mean to get the arithmetic mean of size (1,80,80) as shown in Figure 2(b). These arithmetic mean tiles are stacked up to get a two-dimensional image of the same size as the original, shown in Figure 2(c). Then we subtract the 2D array of Figure 2(c) from the original 2D array of Figure 2(a) to get the resultant image removing the grid artifact shown in Figure 2(d). Then, we check if this process works for all the values along the spectral dimension. It works for all the *m/z* values. After removing the grid artifact for each *m/z* value, we stack all the resultant 2D images to get the final three-dimensional ToF-SIMS dataset used in the NMF and CCA stages.

### 2.4 Co-registration

After performing Sparse NMF, the common tissue section between the three modalities is co-registered using an affine transformation model trained in Pytorch. Preliminary results show good registration between slices from different modalities. The NMF components of these registered tissue sections correspond to co-occurring peaks in the corresponding modes. They may be related to one or more chemical compounds in the sample. First, we register the FT-ICR data to MALDI-TOF data since MALDI-TOF has a higher resolution than FT-ICR but a lower resolution than ToF-SIMS data spatially. Second, we register the MALDI-TOF data to ToF-SIMS data. We get individual binary masks for all the tissue sections present in the spatial image by using *logical not* operation followed by binary dilations. We use Scipy binary fill-holes operation to perform this. Then, we use the generated masks to get a common tissue section between two modalities of data (ex., FT-ICR and MALDI-ToF). Subsequently, we crop and resize the common tissue section to have the same spatial dimension. Generally, we perform downsampling with anti-aliasing. For example, the MALDI-TOF tissue section, with a spatial dimension of 700 *×* 975 pixels, is downsampled to match the corresponding FT-ICR tissue section with a spatial resolution of 65 *×* 97 pixels. To register the common tissue sections, we train a Rigid transformation model in Pytorch and get the rotation and translation matrices in a differentiable way. We use affine grid and grid sample methods from Pytorch to compute the rigid transformation output, which is then used against the reference input to calculate the loss. The loss is then propagated backward to update the parameters in the transformation matrix. After training, we use the trained transformation matrix to register the tissue section with the reference tissue section.

The result of this co-registration process is shown in Figure S7 where an FT-ICR reference tissue section is used to co-register a MALDI-TOF tissue section. Also, we show the downsampled and original MALDI-TOF tissue section after registration with their spatial dimension along with the reference FT-ICR tissue section in Figure S8. We are showing only one NMF component out of the 40 components here.

The MALDI-TOF data is used similarly in the co-registration process to register the ToF-SIMS data. Figure S9 shows the co-registration of a ToF-SIMS tissue section with a MALDI-TOF reference tissue section.

### 2.5 Canonical Correlation Analysis (CCA)

Canonical Correlation Analysis (CCA) is a statistical tool for examining associations between two sets of variables by identifying linear combinations known as canonical variables that maximize their correlation. In the context of this study, the Scikit-learn library’s CCA module was employed, utilizing default settings and specifying 40 components.

Following data registration, CCA was leveraged to discern the physical relationships among various MSI modalities, enabling the prediction of low-resolution FT-ICR MSI images into high-resolution ToF-SIMS. The process unfolded in two stages: initially, CCA was applied to FT-ICR and MALDI-TOF data, facilitating the prediction of low-resolution FT-ICR tissue sections to MALDI-TOF resolution. Subsequently, in the second step, CCA was once again employed, this time with TOF-SIMS data, utilizing the predicted FT-ICR maps obtained in the preceding step to achieve spatial resolution at TOF-SIMS resolution. This iterative approach generated datasets characterized by high spatial and high spectral information. Through these sequential CCA applications, the study aimed to unravel intricate relationships between different MSI modalities and enhance the overall resolution of the acquired data.

## 3 Results

The result of the first step of CCA is shown in Figure 3 where Figure 3(a), Figure 3(c), and Figure 3(e) show the first, second and fifteenth NMF component out of forty of the original FT-ICR data. The second column Figure 3(b), Figure 3(d), and Figure 3(f) show the corresponding NMF component of the predicted FT-ICR data at MALDI-TOF resolution. Here, we can see that the CCA captured the physical relationship between FT-ICR and MALDI-TOF and correctly predicted the NMF components at high resolution. The same colorbar is used to plot these components. The predicted NMF components of all three components have the same intensity profile as the original NMF components.

**Figure 3.**
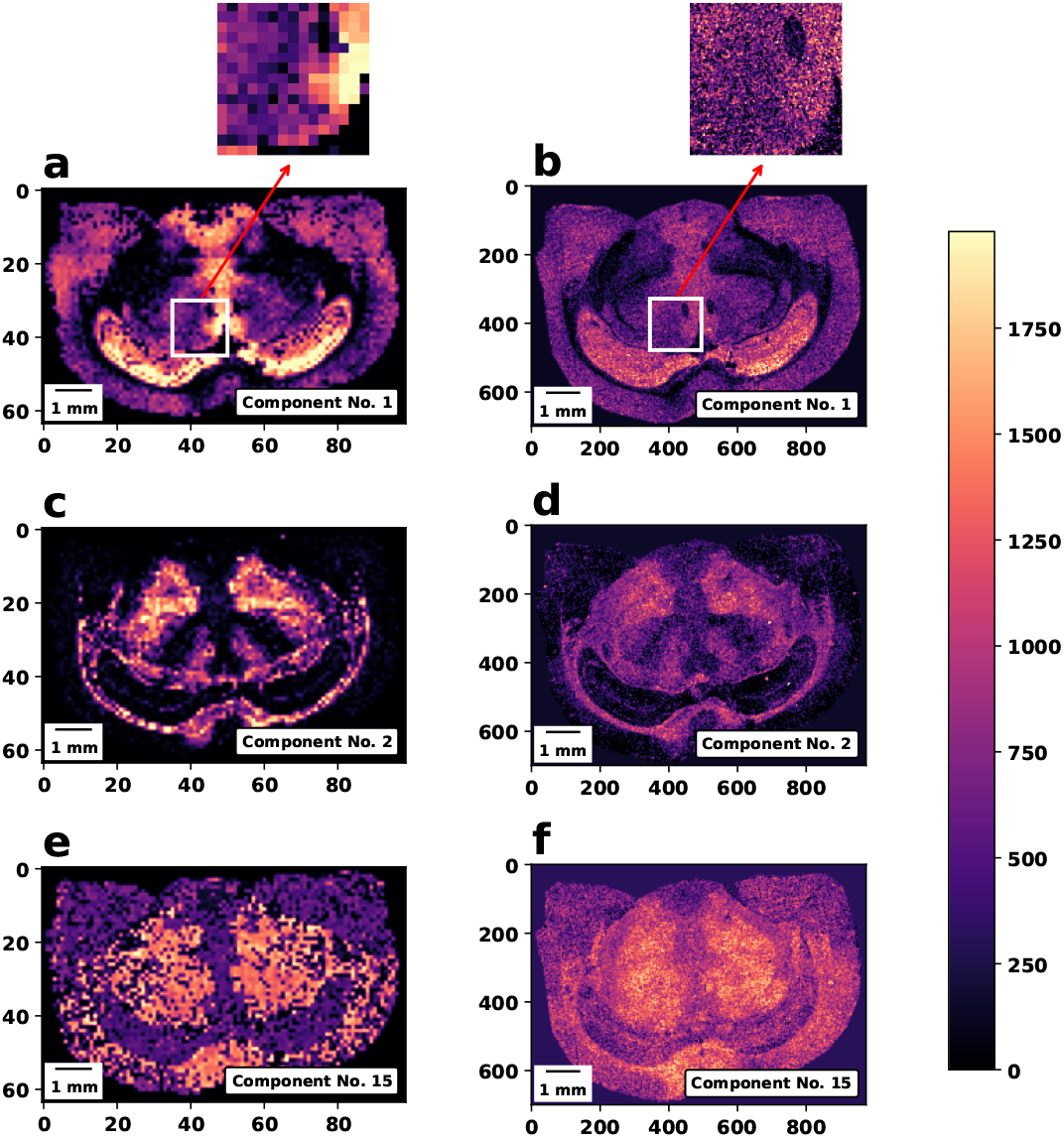
Predicted FT-ICR at MALDI-TOF resolution using CCA. **(a), (c),** and **(e)** show three original FT-ICR NMF components. **(b), (d)**, and **(f)** show the corresponding components of the predicted FT-ICR at MALDI-TOF resolution. A zoomed-in section from original and predicted components are shown on top to show improved features.

In the second step, we use the predicted NMF components at MALDI-TOF resolution, register them with the ToF-SIMS NMF components, and then run the CCA algorithm. The result of the second step of CCA is shown in Figure 4 where Figure 4(a), Figure 4(c), and Figure 4(e) show the second, eighth and seventeenth NMF component out of forty of the original NMF components at MALDI-TOF resolution. The second column Figure 4(b), Figure 4(d), and Figure 4(f) show the corresponding predicted NMF component at ToF-SIMS resolution. Again, it is visible here that the CCA correctly captured the physical relationship between these two modalities of data and predicted the NMF components at high resolution. The same colorbar is used to plot these components. The predicted NMF components of all three components seemingly captured the intensity profile as the original NMF components correctly.

**Figure 4.**
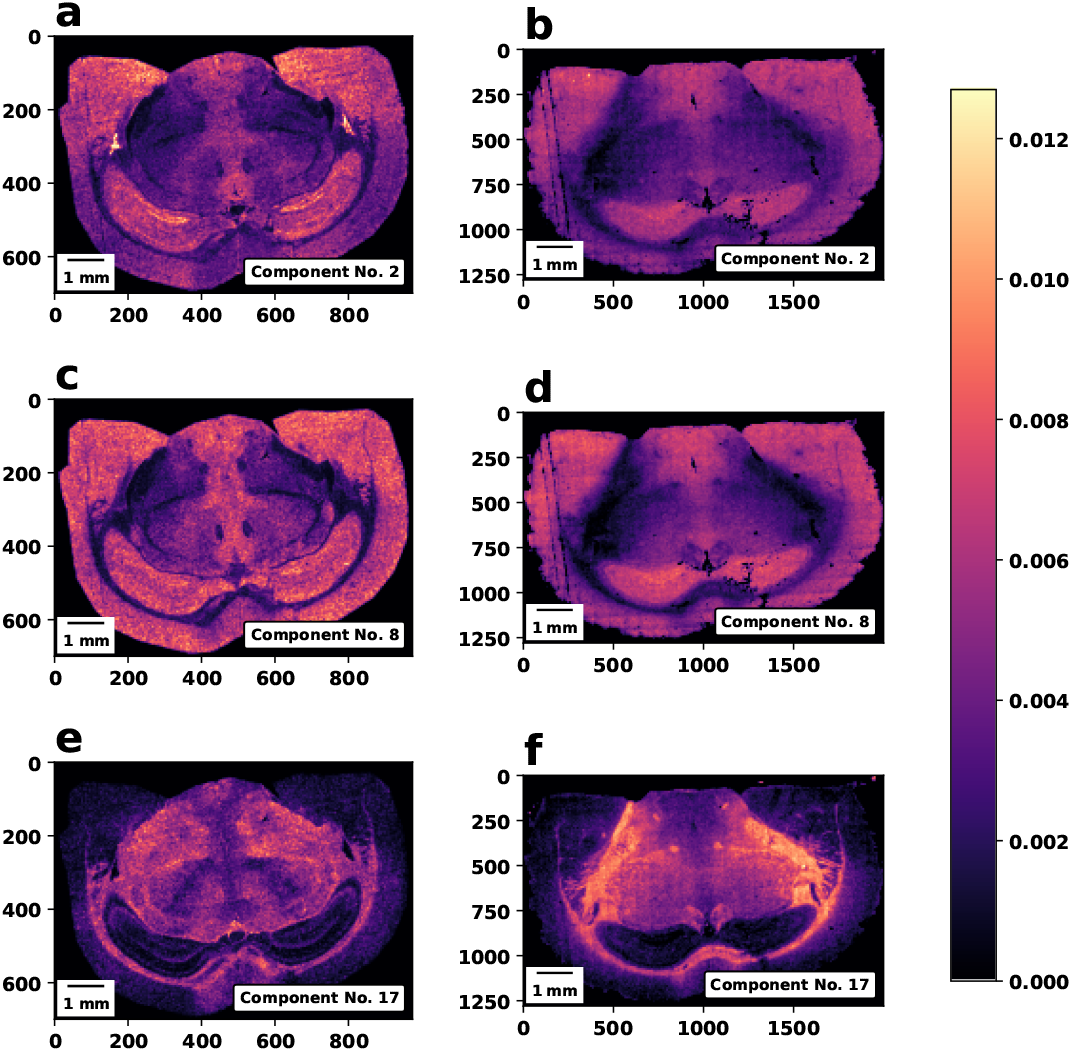
Prediction at ToF-SIMS resolution using CCA. **(a), (c),** and **(e)** show three original NMF components at MALDI-TOF resolution. **(b), (d)**, and **(f)** show the corresponding components of the predicted NMF components at ToF-SIMS resolution.

Moreover, the CCA results are tested against *m/z* peaks, and spectral estimates for the predictions are also observed. Figure 5 shows the 790.8273 *m/z* map for the original FT-ICR data and predicted data at MALDI-TOF and ToF-SIMS resolution. Here, *m/z* values of 790.8273 ± 5 is used to get the mean images as shown in Figure 5(a), Figure 5(b), and Figure 5(c). Again, we can see a similar intensity profile among the three modalities. Also, the mean spectral estimates for a certain area from all three modalities (shown in green for FT-ICR, red for MALDI-ToF, and blue for ToF-SIMS) are calculated and plotted in Figure 5(d), Figure 5(e), and Figure 5(f). Although there is a slight intensity mismatch in the spectral estimates, the peaks are seen to be correctly predicted for both MALDI-TOF and ToF-SIMS prediction. This shows the potential to reveal intricate molecular distributions at high spatial and spectral resolution.

**Figure 5.**
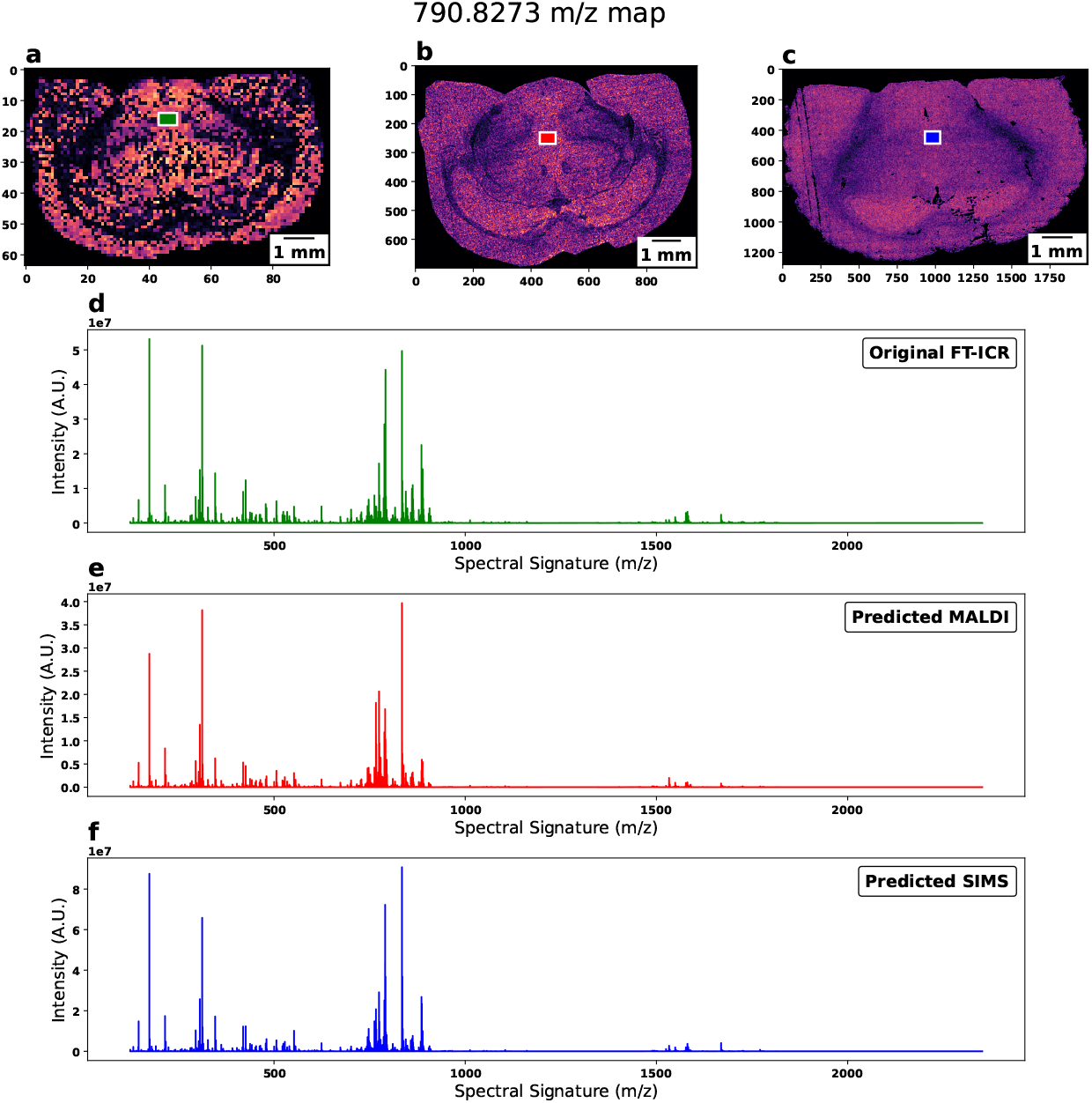
Peak prediction with the predicted data at *m/z* 790.8273. **(a), (b),** and **(c)** show the original FT-ICR, predicted at MALDI-TOF resolution and predicted at ToF-SIMS resolution, respectively. **(d), (e)**, and **(f)** shows the corresponding mean spectral estimate for the three modalities computed from the rectangular box colored in green, red, and blue.

## 4 Conclusion

This investigation showcases the power of mass spectrometry imaging data fusion, achieving submicron spatial and high spectral resolution for complex biological samples. Combining FT-ICR, MALDI-ToF, and ToF-SIMS modalities with advanced image processing and machine learning techniques like CCA yields outstanding results, especially in mouse brain analysis. Future directions include expanding to more tissue sections and creating complete 3D reconstructions of entire mouse brains, promising breakthroughs in neuroscience and beyond. The methodology’s flexibility allows for the seamless integration of additional imaging channels, enabling comprehensive multimodal investigations crucial for more profound insights across scientific domains. Moreover, the adaptability of this approach opens avenues for collaboration across disciplines, fostering interdisciplinary research efforts that push the boundaries of our understanding of complex systems. As this study paves the way for a more holistic comprehension of biological and chemical landscapes, it catalyzes transformative advances with far-reaching implications for materials science, pharmaceutical research, and beyond. Thus, this work reveals immediate discoveries and propels us toward a future where precision and depth redefine our exploration of the intricate interplay within complex systems.

## Supporting information

Supplemental Images

## References

[1] S. Somnath, S. Jesse, G. J. V. Berkel, S. V. Kalinin, and O. S. Ovchinnikova, “Improved spatial resolution for spot sampling in thermal desorption atomic force microscopy mass spectrometry via rapid heating functions,” Nanoscale, vol. 9, no. 17, pp. 5708–5717, 2017. Publisher: Royal Society of Chemistry.

[2] O. S. Ovchinnikova, T. Tai, V. Bocharova, M. B. Okatan, A. Belianinov, V. Kertesz, S. Jesse, and G. J. Van Berkel, “Co-registered Topographical, Band Excitation Nanomechanical, and Mass Spectral Imaging Using a Combined Atomic Force Microscopy/Mass Spectrometry Platform,” ACS Nano, vol. 9, pp. 4260–4269, Apr. 2015. Publisher: American Chemical Society.

[3] R. Van de Plas, J. Yang, J. Spraggins, and R. M. Caprioli, “Image fusion of mass spectrometry and microscopy: a multimodality paradigm for molecular tissue mapping,” Nature Methods, vol. 12, pp. 366–372, Apr. 2015. Number: 4 Publisher: Nature Publishing Group.

[4] A. Belianinov, A. V. Ievlev, M. Lorenz, N. Borodinov, B. Doughty, S. V. Kalinin, F. M. Fernndez, and O. S. Ovchinnikova, “Correlated Materials Characterization via Multimodal Chemical and Functional Imaging,” ACS Nano, vol. 12, pp. 11798–11818, Dec. 2018. Publisher: American Chemical Society.

[5] T. Milillo, R. Hard, B. Yatzor, M. E. Miller, and J. Gardella, “Image fusion combining SEM and ToF-SIMS images,” Surface and Interface Analysis, vol. 47, no. 3, pp. 371–376, 2015. eprint: https://onlinelibrary.wiley.com/doi/pdf/10.1002/sia.5719.

[6] K. upkov, V. Terzopoulos, S. Jain, D. Smeets, and R. M. A. Heeren, “A patch-based super resolution algorithm for improving image resolution in clinical mass spectrometry,” Scientific Reports, vol. 9, p. 2915,Feb. 2019. Number: 1 Publisher: Nature Publishing Group.

[7] J.-H. Rabe, D. A. Sammour, S. Schulz, B. Munteanu, M. Ott, K. Ochs, P. Hohenberger, A. Marx, M. Platten, C. A. Opitz, D. S. Ory, and C. Hopf, “Fourier Transform Infrared Microscopy Enables Guidance of Automated Mass Spectrometry Imaging to Predefined Tissue Morphologies,” Scientific Reports, vol. 8, p. 313, Jan. 2018. Number: 1 Publisher: Nature Publishing Group.

[8] A. Cassese, S. R. Ellis, N. Ogrinc Potonik, E. Burgermeister, M. Ebert, A. Walch, A. M. J. M. van den Maagdenberg, L. A. McDonnell, R. M. A. Heeren, and B. Balluff, “Spatial Autocorrelation in Mass Spectrometry Imaging,” Analytical Chemistry, vol. 88, pp. 5871–5878, June 2016. Publisher: American Chemical Society.

[9] F. Vollnhals, J.-N. Audinot, T. Wirtz, M. Mercier-Bonin, I. Fourquaux, B. Schroeppel, U. Kraushaar, V. Lev-Ram, M. H. Ellisman, and S. Eswara, “Correlative Microscopy Combining Secondary Ion Mass Spectrometry and Electron Microscopy: Comparison of IntensityHueSaturation and Laplacian Pyramid Methods for Image Fusion,” Analytical Chemistry, vol. 89, pp. 10702–10710, Oct. 2017. Publisher: American Chemical Society.

[10] N. Borodinov, M. Lorenz, S. T. King, A. V. Ievlev, and O. S. Ovchinnikova, “Toward nanoscale molecular mass spectrometry imaging via physically constrained machine learning on co-registered multimodal data,” npj Computational Materials, vol. 6, pp. 1–8, June 2020. Number: 1 Publisher: Nature Publishing Group.

