## Supplemental Images for "Predicting High-Resolution Spatial and Spectral Features in Mass Spectrometry Imaging with Machine Learning and Multimodal Data Fusion"

This document provides supporting information for the paper.

### 1 Supplementary Figures

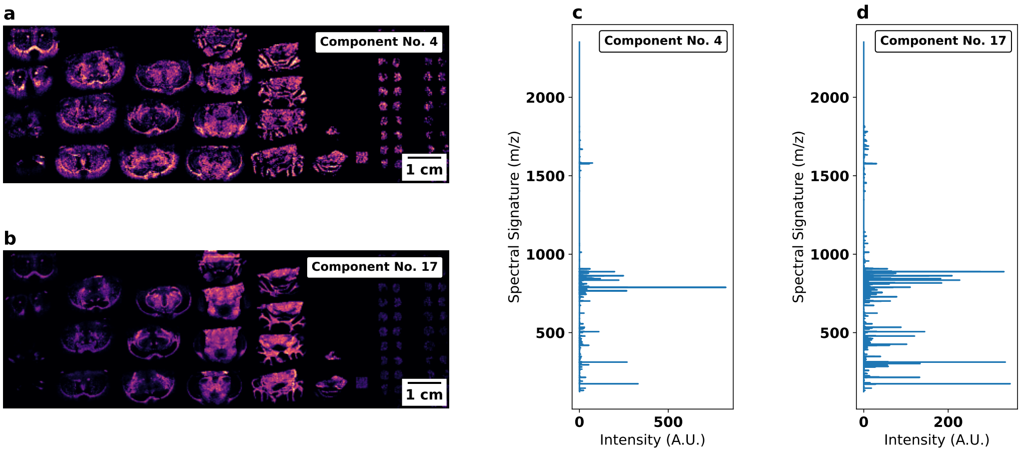


Figure S1: **NMF calculated components for FT-ICR data**. The second NMF component ($W_2$) is laid out spatially in **(a)**, with its spectral counterpart ($H_2$) plotted in **(c)**. The $W_2$ component plotted had 72078 components subsequently mapped to the unique $(x, y)$ location obtained through the sparse NMF process. Similarly, the eighth NMF component ($W_8$) is laid out spatially in **(b)**, with its spectral counterpart ($H_8$) plotted in **(d)**. Visual inspection of the maps generated from the different components demonstrates that different regions of the mouse brain light up in different components, demonstrating that this approach can pinpoint different chemical signatures.

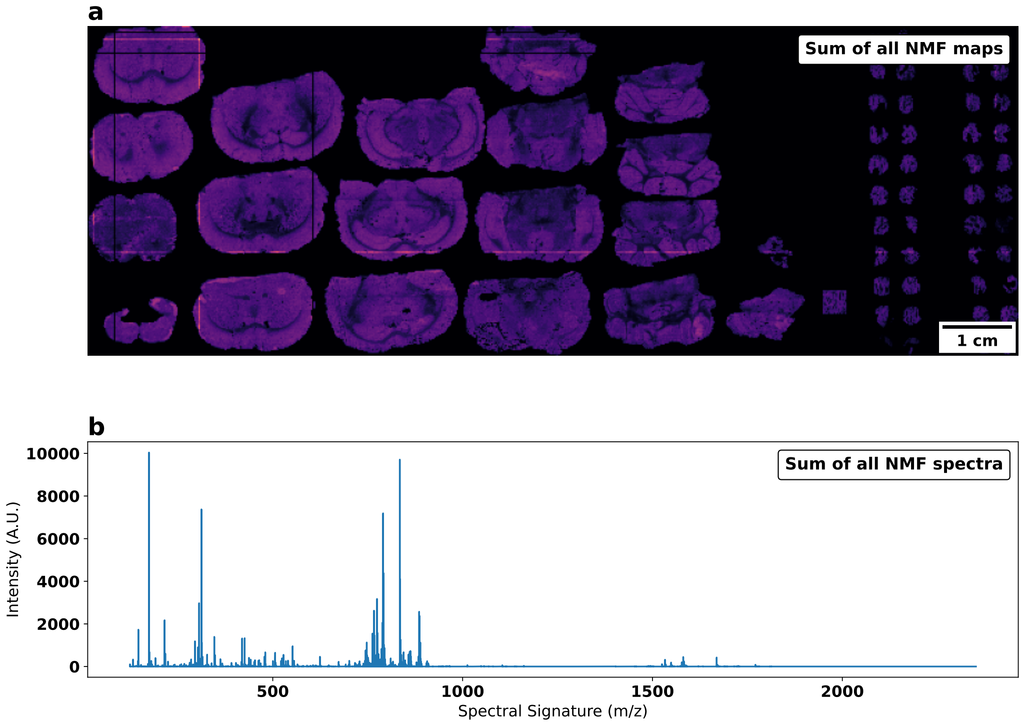


Figure S2: **Sum of solved NMF components for FT-ICR data**. The sum of 40 calculated NMF spatial components is shown in **(a)**, which is very similar to the original spatial map shown in Figure 1(a). Similarly, summing up the spectral maps along its 40 components generated the summed spectra, as seen in **(b)**.

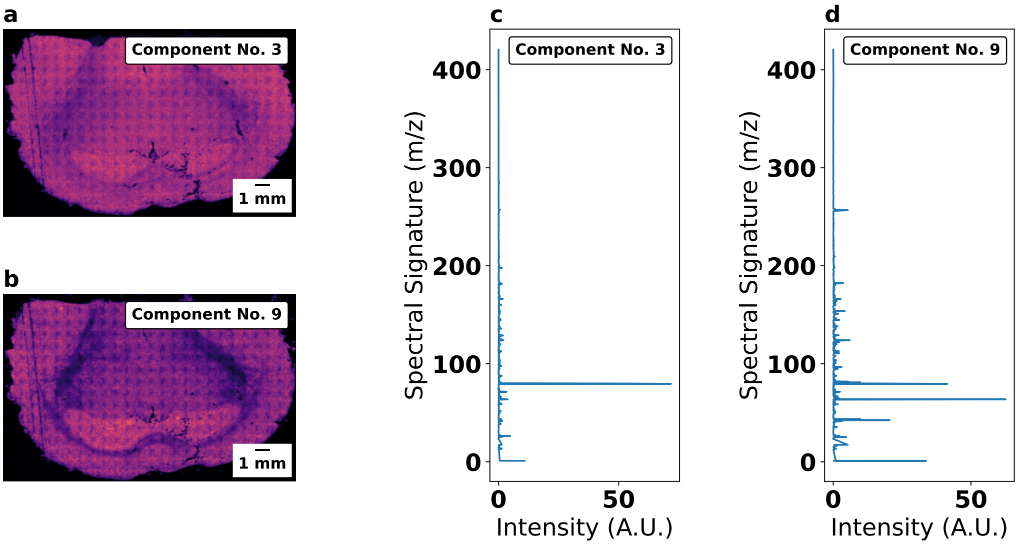


Figure S3: **NMF calculated components for ToF-SIMS data**. The first NMF component ($W_1$) is laid out spatially in **(a)**, with its spectral counterpart ($H_1$) plotted in **(c)**. The $W_1$ component plotted had 2559964 components. The eighth NMF component ($W_8$) is laid out spatially in **(b)**, with its spectral counterpart ($H_8$) plotted in **(d)**.

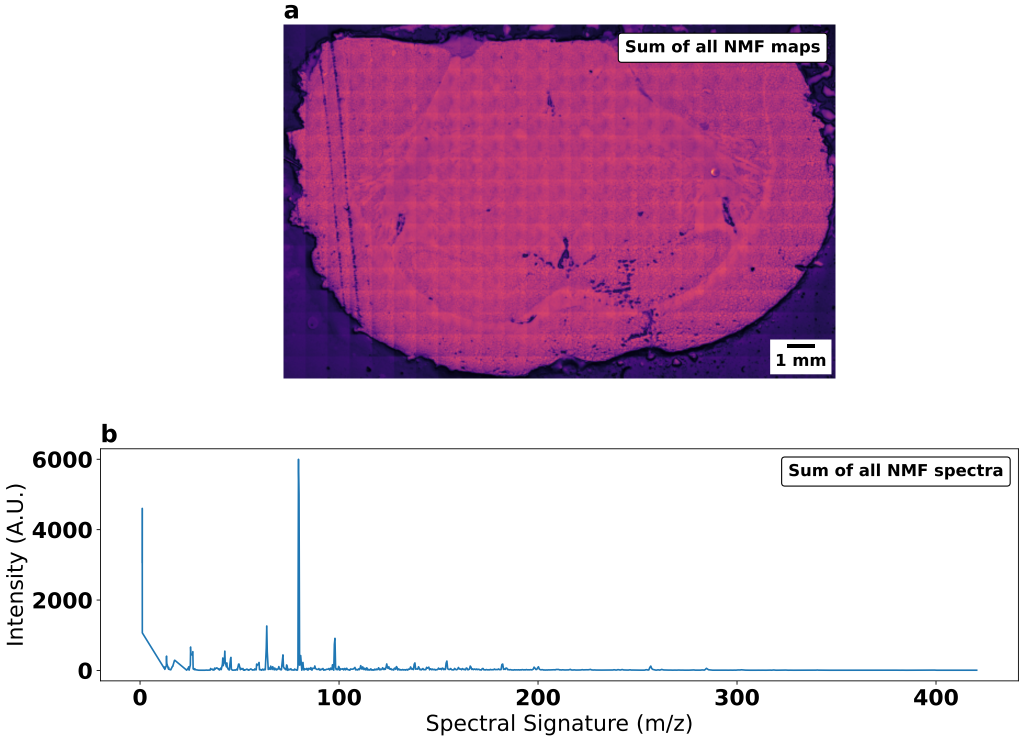


Figure S4: **Sum of solved NMF components for ToF-SIMS data**. **(a)**, Sum of 40 calculated NMF spatial components. **(b)**, Sum of 40 NMF spectral components.

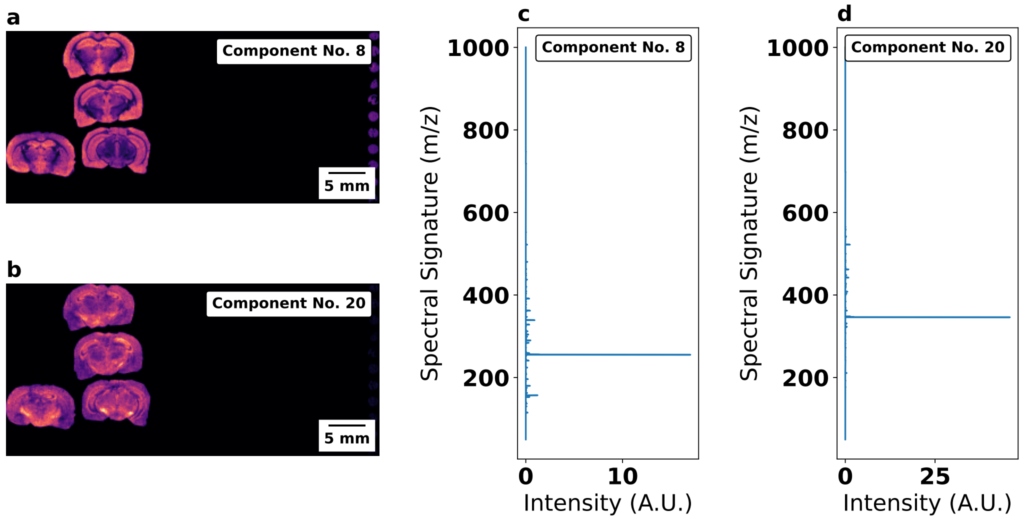


Figure S5: **NMF calculated components for MALDI data**. The first NMF component ($W_1$) is laid out spatially in **(a)**, with its spectral counterpart ($H_1$) plotted in **(c)**. The $W_1$ component plotted had 2030712 components. The eighth NMF component ($W_8$) is laid out spatially in **(b)**, with its spectral counterpart ($H_8$) plotted in **(d)**.

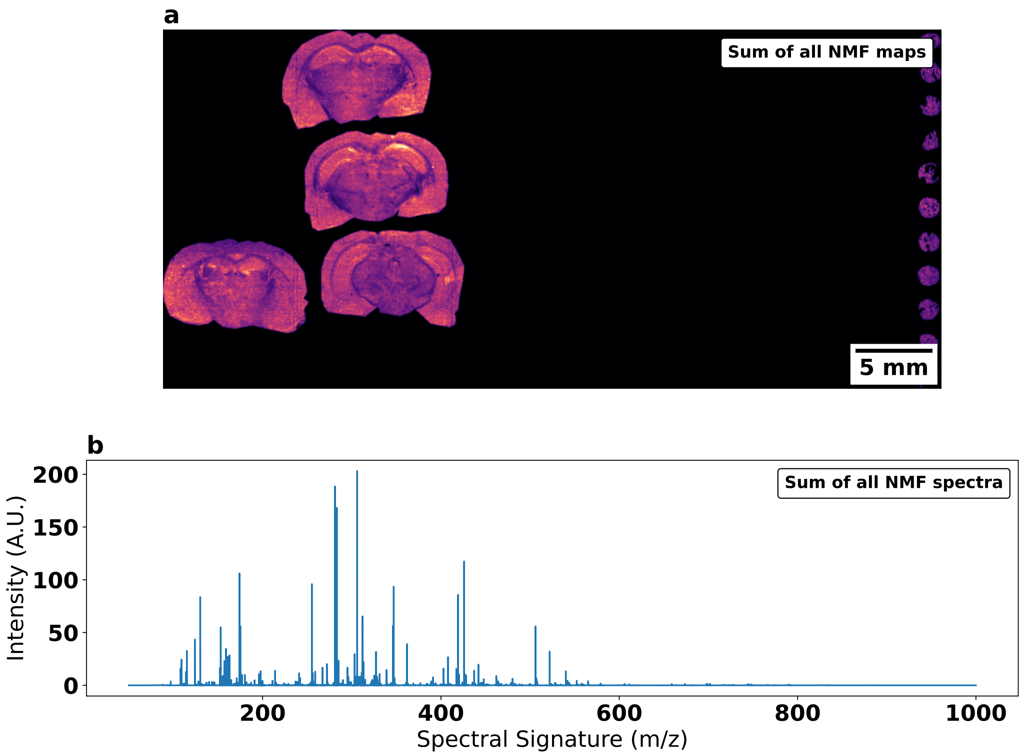


Figure S6: **Sum of solved NMF components for MALDI data**. **(a)**, Sum of 40 calculated NMF spatial components. **(b)**, Sum of 40 NMF spectral components.

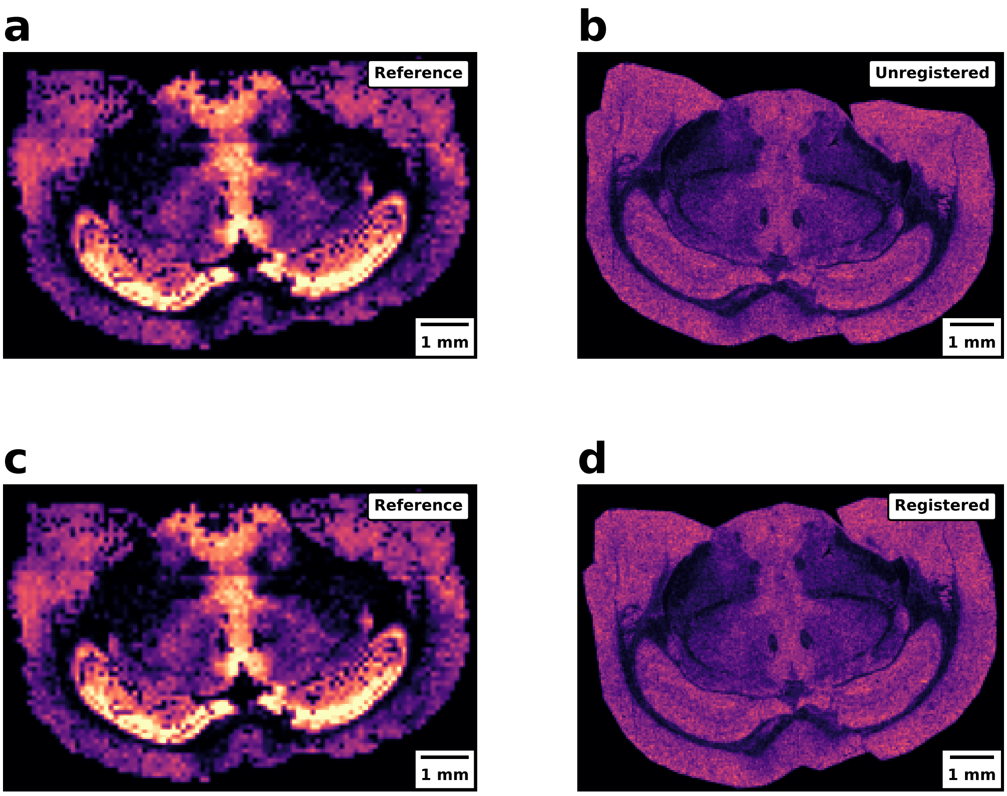


Figure S7: **Co-registration of MALDI to FT-ICR**. **(a)**, and **(c)** Reference FT-ICR tissue section. **(b)** Unregistered MALDI tissue section. **(d)** Registered MALDI tissue section.

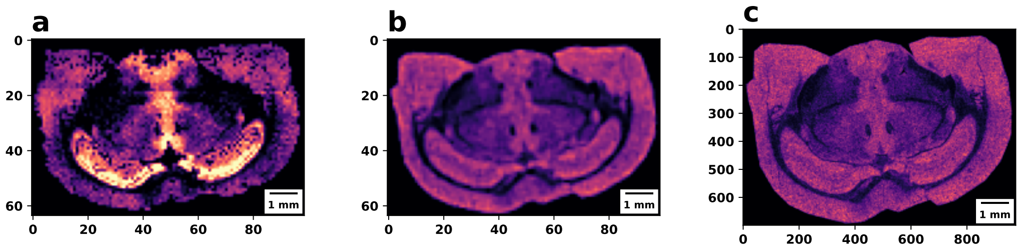


Figure S8: **Downsampling MALDI data after registration with spatial dimension**. **(a)** reference FT-ICR tissue section, **(b)** Resized MALDI tissue section after registration, **(c)** original MALDI tissue section after registration.

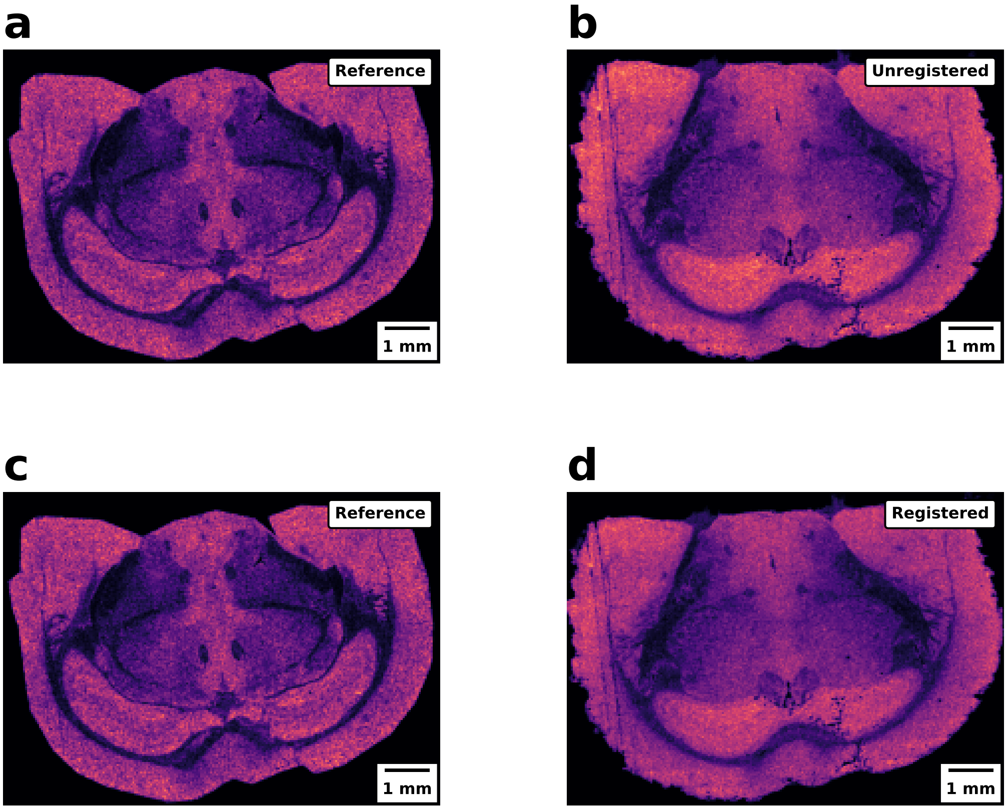


Figure S9: **Co-registration of ToF-SIMS to MALDI**. **(a)**, and **(c)** Reference MALDI tissue section. **(b)** Unregistered ToF-SIMS tissue section. **(d)** Registered ToF-SIMS tissue section.
